# Impact of overfishing on density and test-diameter size of the sea urchin *Tripneustes gratilla* at Wakatobi Archipelago, south-eastern Sulawesi, Indonesia

**DOI:** 10.1101/727271

**Authors:** La Nane

## Abstract

Sea urchin *Tripneustes gratilla* is one of an economic important fisheries resources product for localities at Wakatobi archipelago. High demands for sea urchin gonad have intensified high fishing activity. We hypothesize that sea urchins at Wakatobi have overfished. To answer that hypothesizes; we measure the density and its test diameter size at two different sites. They are Pulau Tomia (inhabited area) and Pulau Sawa (uninhabited area and very distant from the localities). The results show that sea urchin density and its test diameter is significantly different. The densities (mean ± SE) Tripneustes gratilla at Pulau Sawa and Pulau Tomia are 10 ± 0.6 (ind./m^2^) and 2.7 ± 0.9 ind./m^2^, respectively. Moreover, the test diameter at Pulau Sawa and Pulau Tomia are 69.7 ± 2.1 mm (mean ± SE), and 58.5 ± 1.7 mm (mean ± SE), respectively. These results have indeed shown that overfishing has occurred. Therefore, sea urchin with test diameter 66–75 mm, 76–85 mm, and 86–95 mm have disappeared at Pulau Tomia. Our conclusion reveals that fishing of sea urchin Tripneustes gratilla at Pulau Tomia has overfished.

## 1. Introduction

Sea urchins have been known as major ecologically important components, especially in shallow tropical seas (Steneck, 2013; Luja & Malay, 2019). They also have played a crucial rule in nutrient cycling, macroalgal grazing, and bioerosion (Koike et al. 1987; Bronstein and Loya 2014; Ling et al. 2018). The roe (gonad) of sea urchins have been utilized as seafood products in many parts of the world, particularly Asia and Europe (Baiao et al., 2019; Kato, 1972; Salvo et al., 2016; Taylor et al. 2017). Thereby, Sea urchin gonads have highly valued commercially (Lawrence, 2007; Brown and Eddy, 2015; Sun & Chiang, 2015). As a consequence, demand on sea urchin gonads is rapidly increasing globally (Brown and Eddy 2015, Mos & Dworjanin, 2019). But that high demands have eased the capture of sea urchin and the long recruitment time have also caused overfishing to wild populations of sea urchin (Sloan 1985; Keesing & Hall 1998; Lesser & Walker 1998; Andrews et al. 2002; Robinson 2004).

In Wakatobi archipalego, Sea urchin *Tripneustes gratilla* is also one of an economic important fisheries resources product that has been fished traditionally for commercial purpose for the years. However, globalization that followed with a high birth-rate of human at Wakatobi, demand for sea urchin gonad has also increased. As a consequence, the number of fished sea urchin was also intensified and reduced the population size in nature. Therefore, the number of individuals at fishing ground seems to be low in number and to be small in test-diameter size.

Sea urchins as the natural resources could be overexploited by high fishing pressure. Decreasing sea urchin at nature may be related to the fishing activity that has been intensified since the last decade. Since fishing activity to catch sea urchin has also escalated the number of fishing gears and fishing boats. As a result, intense fishing trips to catch sea urchin have also increased. In parallel with that, the total harvest to sea urchins is also large in quantities.

On the other hand, sea urchin stock is not only limited in number but also decreased on test-diameter size. Moreover, the fishing ground is also to be distant from the localities. Thus, Fishers need more effort and cost to catch sea urchin. Those indications are very strongly indicated the overexploitation. According to Widodo and Suadi [as cited in Dewi et al. 2019], the characteristics of overfished are the time taken to go out to sea being longer, the fishing locations tending to be distant, the productivity or catch per unit effort tending to decrease, the size of the target fish is getting smaller, and the cost of the catching operations increasing.

Therefore, for answering the hypothesis; are sea urchins *Tripneustes gratilla* have overexploited at Wakatobi. We performed this study to verify and measure the density and its test diameter size decreasing. Thereby, this study is very important as part of an effort to manage the sea urchin resources, especially the sea urchin *Tripneustes gratilla*.

## 2. Material and methods

### 2.1 Study site

This study was conducted on August 2018 at Pulau Tomia (38° 8’U, 141° 57’T) and Pulau Sawa (38° 8’U, 141° 57’T), Wakatobi Regency, South eastern Sulawesi, Indonesia (see Figure 1). Sea water depth at both sites is 2 m depth at a high-tide level, and 0.01 m at a low-tide level. Sea bottom substrate was dominated by seagrass beds with sandy substrate. The samples measurement was performed in the field and conducted when the seawater retreat at lower tide level. Both Pulau Sawa and Pulau Tomia was considered as the study sites due to the fishing intensity of fishermen at both sites.

**Figure 1.**
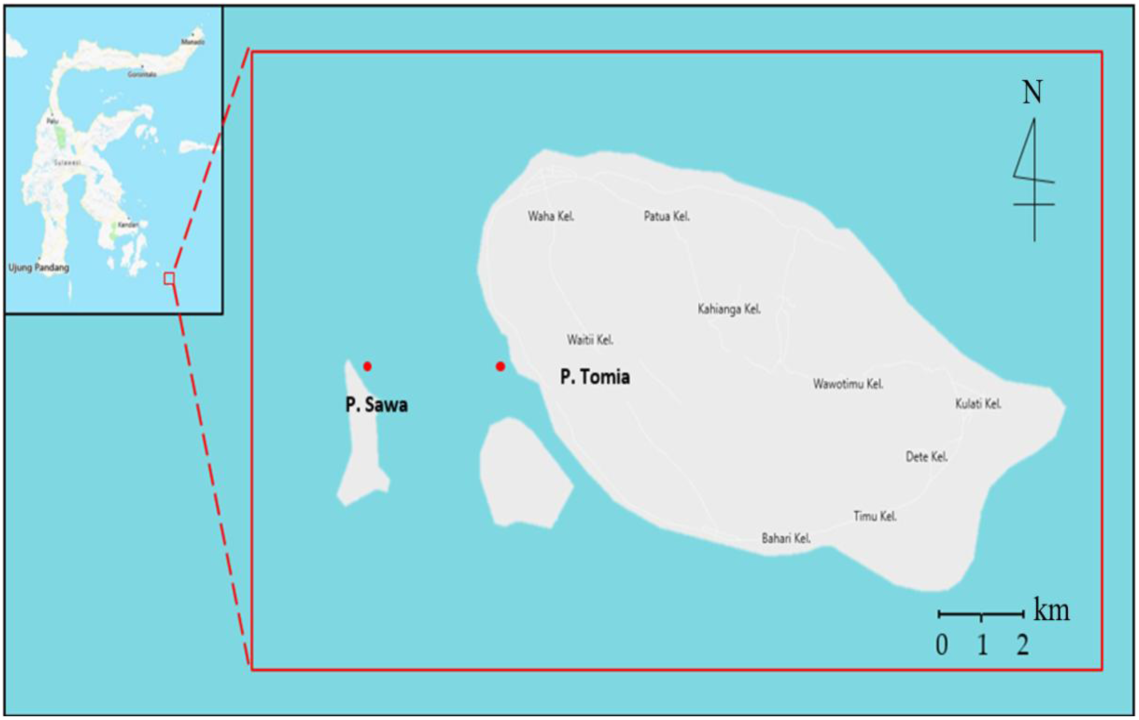
Study area. The red dot (•) indicate the coordinate position of both study sites.

### 2.2 Measurement

#### 2.2.1 Density (ind./m^2^)

Sea urchin density at both sites was randomly collected using a 1 m × 1 m transect quadrate with three replication along the seagrass beds when the seawater is retreading at the lowest level (approx. 0.01 m in depth). Then, the number of sea urchin per quadrate was counted as density and noted on aqua-note.

#### 2.2.2 Test diameter (mm)

In parallel with the density collection and measurement, each individual of sea urchin was measured with a caliper (accuracy 0.1 mm) and noted.

#### 2.2.3 Statistical analyses

All statistical analyses were performed using SPSS Statistic IBM 22^®^ (SPSS Inc., Chicago, IL, USA). Significant difference among sea urchin density (ind./m^2^) and its test diameter (mm) between Pulau Sawa and Pulau Tomia were analyzed using *t-*test where the significant difference is *p* < 0.05.

## 3. Results

### 3.1 Density (ind./m^*2*^)

There is a significant difference in sea urchin density betwen Pulau Sawa and Pulau Tomia (*p* < 0. 05). The densities (mean ± SE) of the sea urchin *Tripneustes gratilla* at Sawa Pulau and Tomia Pulau are 10 ± 0.6 ind./m^2^, and 2.7 ± 0.9 ind./m^2^, respectively (see Fig. 2).

**Figure 2.**
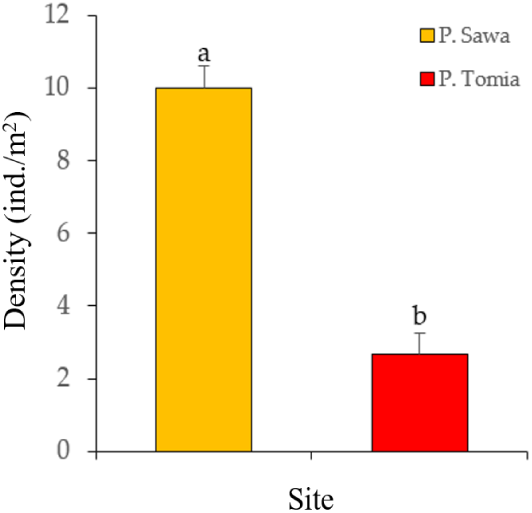
Densities (mean ± SE) of of sea urchin *Tripneustes gratilla* at both sites; where(F (0.7) = 0.44, *p* = 0.02.)

### 3.2 Test diameter (mm)

There is a significant difference in test diameter sea urchin *Tripneustes gratilla* at both sites. Test diameter (mean ± SE) at Pulau Sawa is 69.7 ± 2.1 mm, and Pulau Tomia is 58.5 ± 1.7 mm (see Fig. 3)

**Figure 3.**
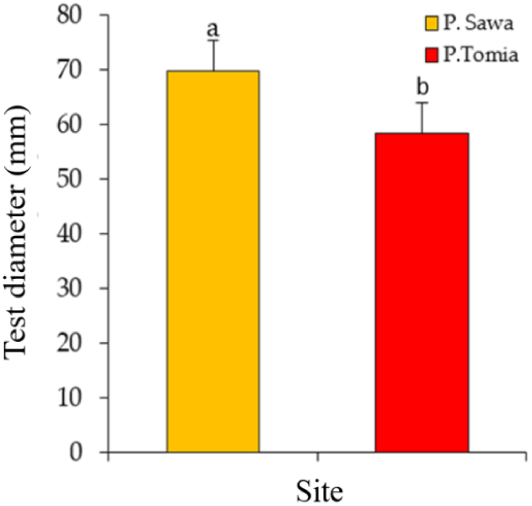
Test diameter difference (mean ± SE) of the sea urchin *Tripneustes gratilla* among Pulau Sawa and Pulau Tomia. Different letter on bar indicate signifikat differences (*p* < 0.05).

### 3.3 Test diameter size frequency

Distribution frequency of sea urchin test-diameter at both sites are different (see Fig. 4). Test diameter with a large size at pulau Tomia and Pulau Sawa is different in number. The larger sea urchin at Pulau Sawa has dissappered.

**Figure 4.**
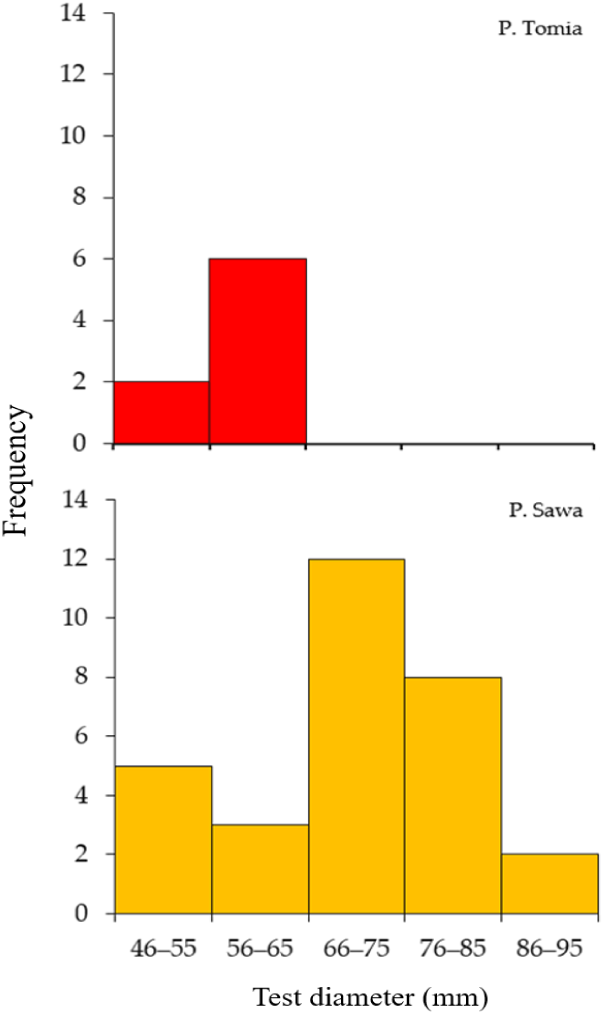
Differences of distribution ferequency of sea urchin test diameter *Tripneustes gratilla* between Pulau Sawa and Pulau Tomia.

## 4. Discussion

### 4.1 Density

Decreasing of sea urchin density at Pulau Tomia (inhabited area) and Pulau Sawa (uninhabited area) show that fishing pressure positively occurred and it significantly reduced the number of sea urchins. The distance of fishing ground with fishers’ resident seems to have a strong impact on that fishing pressure. It can be proved by the significant decreasing of sea urchin number at Pulau Tomia compare to the sea urchin density at Pulau Sawa. These findings indicate that overfishing is truly happening. As a consequence, the urchin’s number cannot recover immediately.

### 4.2 Test diameter

According to Widodo and Suadi (as cited in Dewi et al., 2019), the indication of overfishing can be identified by decreasing fishing size. In parallel with these arguments, our finding also shows significant differences in the distribution of sea urchin test. The average distribution of test diameter at Pulau Sawa is higher than the average of sea urchin test diameter at Pulau Tomia.

### 4.3 Distribution frequency of sea urchin test diameter

The disappearance of sea urchin with test diameter size 66–75 mm, 76–85 mm, and 86–95 mm at Pulau Tomia has revealed that sea urchins *Tripneustes gratilla* have overfished. The lost of that big sea urchin size may have affected by fishing activity that have occurred for the years. As a consequence, sea urchin cannot recover immediately due to high fishing pressure that still happened.

## 5. Conclusion

Decreasing of sea urchin number (density) and sea urchin test-diameter at Pulau Tomia that distant with fisher’s resident than sea urchin at Pulau Sawa (unresidented area) indicate that fishing activity was over. As a consequence, the size of sea urchin at Pulau Tomia is smaller than sea urchin test diameter at Pulau Sawa due to recovery capability.

## Acknowledgement

Thanks to Mr. Dian Sardin for help during the collection of sea urchin. I also thank to Mrs. Nurqadri Syaia Bakti for the technical support during this study.

